# Increased brain growth in escaped rainbow trout

**DOI:** 10.1101/2021.06.17.448828

**Authors:** Frédéric Laberge, Marie Gutgesell, Kevin S. McCann

## Abstract

Recent examples of rapid brain size plasticity in response to novel laboratory environments suggest that fish brain size is a flexible trait, allowing growth or shrinkage of brain tissue based on short term needs. Nevertheless, it remains to be seen if plasticity of fish brain size is relevant to natural environmental conditions. Here, using rainbow trout escaped from a farming operation as a natural experiment, we demonstrate that adult fish brain size can change rapidly in response to life in a natural lake environment. Specifically, escaped trout had on average 15% heavier brains relative to body size than captive trout after living for about 7 months in the lake. Because relative brain size of most escaped trout fell above the range of variation seen within the captive trout population, we conclude that increased brain size was achieved by plasticity after escape. Brain morphology analysis showed that the most anterior regions (olfactory bulbs and rest of telencephalon) contributed most to the increase in overall brain size in escaped trout. Relative size of the heart ventricle, another organ which can be subject to plastic changes under variable environmental conditions in fish, did not differ between escaped and captive trout. Massive and selective brain growth under the changed environmental conditions associated with escape from holding pens highlighted the plastic potential of fish brain size and suggests that a shift to increased complexity of life in the wild setting of a lake imposed greatly increased cognitive requirements on escaped trout.

## Introduction

Living organisms are constantly faced with changing conditions on minute, daily, seasonal to inter-generational time scales. Scientists have argued that ecosystems are prototypical examples of complex adaptive systems with organisms capable of rapidly responding to changing conditions in a manner that fundamentally mediates ecosystem stability and function (Levin, 1998). Following these ideas, evolutionary ecologists have shown that rapid evolutionary responses can mediate ecosystem stability (Yoshida et al., 2003) and food webs ecologists have shown that behavioral foraging responses of highly mobile top predators, if rapid, can also act as potent stabilizers in a noisy world (McCann and Rooney, 2009). It remains unclear if plastic change in physiological systems at a time scale faster than evolutionary change can act in support of such stabilizing behavioral foraging responses, because empirical field research on rapid phenotypic flexibility is limited.

The ability to reversibly change organ systems to match rapid or predictable change in environmental conditions within a lifetime, termed phenotypic flexibility by Piersma and Lindström (1997), has been demonstrated in the digestive system of representative species of all vertebrate groups [mammals (Hammond et al., 2001), birds (Piersma et al., 1993), reptiles (Naya and Bozinovic, 2006; Secor, 2008), amphibians (Naya et al., 2009), fish (Armstrong and Bond, 2013; Blier et al., 2007)]. Before digestion can begin, predatory species rely on a combination of cognitive and locomotor abilities for prey capture; therefore, matching changes in the nervous and cardiovascular systems with ongoing foraging conditions through sufficiently rapid phenotypic flexibility would be adaptive. Fish could maintain a lifelong potential for plasticity of brain size because of widespread adult neurogenesis (Kaslin et al., 2008) and the modulation of neurogenesis and brain size by sensory experience (Hall and Tropepe, 2020). Similarly, the ability to display cardiac remodelling in response to experimental manipulations of temperature (Keen et al., 2017) or exposure to stressors (Johansen et al., 2017; Simonot and Farrell, 2007) suggests that fish hearts are highly plastic. Such features make fish good models to assess the extent of phenotypic flexibility associated with changes in foraging demands. Additionally, comparison of captive and wild fish have shown larger brains (Marchetti and Nevitt, 2003; Mayer et al., 2011; Park et al., 2012) and heart ventricles (Graham and Farrell, 1992) in wild fish, suggesting that life in a natural environment puts important demands on both the nervous and cardiovascular systems, which are met by investment of energy into organ growth and maintenance. Since laboratory experiments cannot completely capture the richness of experience in a natural environment, we sought an opportunity for a natural experiment where captive fish would escape from a floating pen culture operation and forage on wild prey before they were sampled.

Escape from pen culture operations happens regularly and escapees usually establish in the local environment, at least for a short period of time (Charles et al., 2017; Naylor et al., 2005). Patterson and Blanchfield (2013) obtained about 50% survival of marked or tracked rainbow trout 3 months after simulated escape from aquaculture pens in Lake Huron, and recaptured trout up to 2.5 years after release. Most concerns about fish escapes so far have revolved around competition between escaped fish and the local fauna. Here, we used escape from growing pens as an opportunity to study the effects of an abrupt transition from captive to natural environmental conditions on fish organ plasticity. Rainbow trout were sampled approximately 7 months after a large escape event due to a fall storm at a freshwater aquaculture operation located near a long-term sampling site in Lake Huron, Ontario, Canada. Because the escape event happened shortly before harvest (trout approx. 1 kg body mass), the potential effects would be limited to late stages of life. Experimental manipulations entailing environmental enrichment or the transition from a natural environment to captivity have produced changes in relative brain size in adult fish within a period of three to six weeks (Fong et al., 2019; Herczeg et al., 2015; Park et al., 2012; Turschwell and White, 2016). This ability supports the hypothesis that fish maintain the capacity to display phenotypic flexibility of brain size in response to changes in environmental complexity throughout life. If this hypothesis is true, we can predict that the adult-size trout that escaped from pens and foraged in a complex natural lake environment would show an increase in relative brain size compared to trout directly sampled from growing pens. Similarly, increased swimming demands are likely associated with life in a large lake compared to the restricted space available in captivity, which should promote a heart ventricle phenotype adapted for better swimming performance like the larger ventricles seen in wild rainbow trout (Graham and Farrell, 1992). Thus, we also predicted that escaped trout would have larger heart ventricles compared to trout sampled from growing pens to meet increased swimming demands in the lake environment.

## Materials and Methods

### Study system

Escaped trout were collected in Parry Sound, Ontario, Canada. Parry Sound is a large body of water (about 12 × 10 km) connected to Lake Huron’s Georgian Bay by a shallow channel approximately 6 km long, creating a natural barrier to fish population movements. A commercial rainbow trout pen culture operation is located in the southernmost part of Parry Sound in Depot Harbour (Figure 1A). Although self-reproducing populations of rainbow trout and chinook salmon are found in Georgian Bay and some of its tributaries (Dobiesz et al., 2005; Johnson et al., 2010), Parry Sound is distinguished by the exclusive presence of a lake trout population as the top pelagic fish predator (Reid et al., 2001). Lake trout (1981-1997) and rainbow trout (1986-1994) were stocked in Parry sound to subsidize sport fishing (Reid et al., 2001); however, while lake trout achieved successful reintroduction criteria in Parry sound by 1997 and remain abundant (Trumpickas et al., 2020), rainbow trout have not been stocked since 1994 and there is no evidence of natural reproduction of this species in this area.

**Figure 1:**
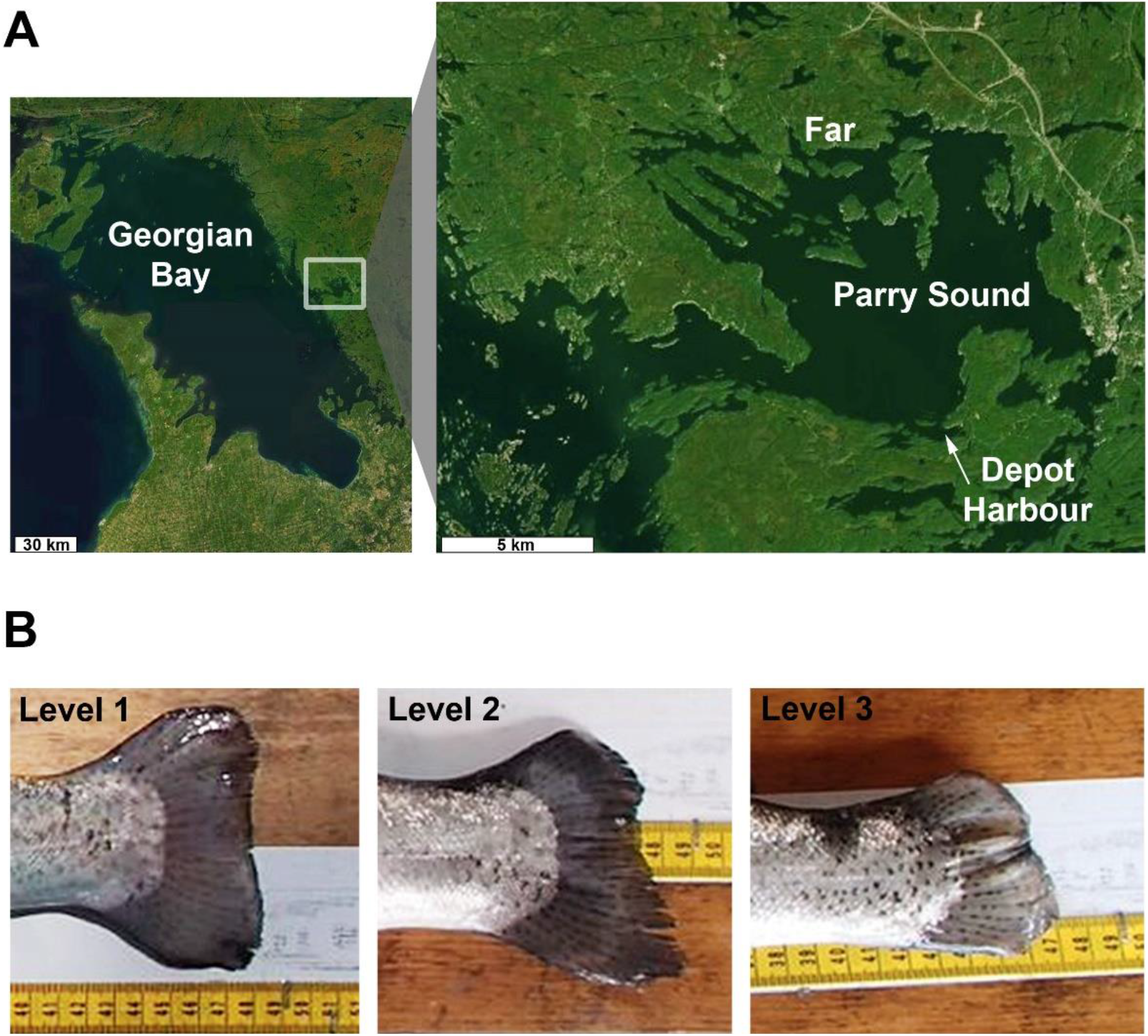
Sampling site and fin erosion of the rainbow trout of Parry Sound, Ontario. A) Parry Sound is a body of water connected to Lake Huron’s Georgian Bay by a channel creating a natural barrier to fish movements. Captive trout were taken from the commercial floating pen culture operation located in Depot Harbour in southern Parry Sound (arrow). Escaped trout were sampled both near Depot Harbour and far away from the pen culture operation in northern Parry Sound (Far). Maps source: Esri World Imagery, Nov. 19, 2020 (www.arcgis.com). Maps generated using R (v. 4.0.3). B) Examples of the three levels of caudal fin erosion in rainbow trout of Parry Sound: level 1 (little to no erosion), level 2 (intermediate) and level 3 (advanced). Little erosion was seen in wild lake trout of Parry Sound while the high incidence of fin erosion in captive fish housed at high density in pens carried over to the escaped rainbow trout sampled in Parry Sound.

### Sampling and preparation

Escaped trout were sampled in the last week of May 2019 (n=26) using angling and gill nets. The fish were sacrificed by a light stunning blow followed by neck puncture to section the spinal cord caudal to the brain. Escape was due to a 2018 fall storm that damaged growing pens and led to the escape of tens of thousands of fish into the lake. A baseline for comparison with escapees relied on sampling of captive trout directly from growing pens in 2019 as well as in prior years (2015-2017), part of a study of resource subsidies of the farming operation into the Parry Sound food web (Johnson et al., 2018). The use of fish sampled over multiple years was needed to establish a reliable baseline of captive fish. Captive fish were collected in June-August 2015-17 (n=5-9 per year), May 2019 (n=10) and early September 2019 (n=6). The September 2019 fish were needed because the only captive trout available in May 2019 were unusually small (mean and range fork length 276 [237-309] mm compared to 417 [357-555] mm in other years). Most captive fish were taken directly from growing pens using dip nets and sacrificed as described above, but the September 2019 collection slightly differed in that the fish taken from the growing pens were transported in iced water to a processing plant before they were obtained for tissue sampling and preparation. Baseline organisms (mayfly larvae, snails, zebra mussels) and feed (2.5mm, 4mm, 5mm, 6mm, and 7.5mm Premium Trout FW pellets, Skretting Inc.) were obtained in May 2019 and summer 2018, respectively. Sampling procedures were approved by the Ontario Ministry of Natural Resources (permits UGLMU2016-06, UGLMU2016-05, UGLMU2017-05 and UGLMU2019-04) and the University of Guelph animal care committee (protocols 3155 and 3563).

Fish were processed daily on shore (or in the lab for the September 2019 captive fish). This involved taking a photograph of the whole fish, weighing body mass to the nearest 0.01 kg with a Rapala Pro Select digital scale, and measuring fork length to the nearest 1 mm on a measuring board. Fish body cavities were opened to examine gonads, obtain a liver tissue sample and remove the heart. A sample was then taken from the dorsal caudal musculature (skin cut out). Remaining skin (including scales) from both sides was separated from muscle using a filleting knife. Then, the top half of the head was dissected, and the base of the braincase exposed and cut gently to allow access to the otoliths. Fine tweezers were used to remove the otoliths without damaging the brain. All samples of liver, muscle, skin and otoliths were frozen at -20 °C immediately and kept frozen until processing for stable isotope analysis (see below). The top half of the head and the heart were immersed in fixative (10% buffered formalin) and remained in this solution until further dissection, which happened every year within a maximum of 8 months after collection. Yearly weighing of the same brain and heart ventricle samples (n=12) at different intervals showed an average 3.5%, 4.7%, and 29% decrease in mass after 1.5, 2.5 and 3.5 years in formalin storage, respectively, suggesting minimal decrease in the few months of storage that preceded data acquisition. Brains were dissected out of the fixed heads, trimmed of excess cranial nerves, and the spinal cord was cut at the level of the obex. Brains were then blotted using Kimwipes (Kimberly-Clark) to remove excess formalin before weighing using an analytical balance (Accu-124D Fisher Scientific) at a resolution of 0.0001 g. Heart ventricles were trimmed of surrounding tissue before blotting and weighing in the same manner. The relationships between fish body weight and organ weights were thus between ‘wet’ body weight and ‘post-fixation’ organ weights.

Brain and heart ventricle morphology were also assessed to determine if these parameters were influenced by escape. Assessment of brain morphology was based on Edmunds et al. (2016). Briefly, the volumes of five brain regions (olfactory bulbs, rest of telencephalon, optic tectum, cerebellum, hypothalamus) were estimated using the ellipsoid method: Volume = π/6 (Length × Width × Height) (White and Brown, 2015). Digital images of the dorsal, ventral and left sides of the brain were taken through an Olympus SZ61 dissection microscope using a Cannon Powershot G9 digital camera and PSREMOTE v.1.7 software. The linear length, width, and height of brain regions were measured using the straight line measuring tool in Fiji ImageJ (Schindelin et al., 2012). Only the left side of the brain was photographed by assuming that the height of both sides of bilaterally symmetrical brain regions was the same. Heart ventricle shape was assessed because an elongated heart ventricle characterizes wild trout (Poppe et al., 2003) and is a phenotype associated with better swimming performance (Claireaux et al., 2005). Digital callipers (Mastercraft) were used to measure maximal length and width of the fixed heart ventricles to the nearest 0.1 mm to obtain a basic measure of shape, the length to width ratio. Length was obtained between the side where bulbus arteriosus and atrium are attached to the ventricle and the posteriorly oriented tip of the pyramid-shaped ventricle. Width was obtained at a right angle to the length measurement between the dorsal and ventral, or lateral, ventricular surfaces, whichever was widest.

### Isotope analysis

Stable isotope ratios of carbon and nitrogen were measured to infer differential resource use between escaped and captive trout (Vander Zanden and Rasmussen, 1999). Tissues with different molecular turnover rates were analyzed as we only expected divergence in isotopic signatures in tissues recently turned over (liver fastest followed by muscle) between escaped and captive trout based on differential consumption of wild prey and trout feed. Common isotopic signatures between escaped and captive trout in tissues with low turnover rates (scales and otoliths) would support feeding on a common resource (trout feed) prior to escape. In preparation for stable isotope analysis, fish liver and muscle samples, baseline organisms and feed were dried at 70 °C for 2 days and ground into a fine powder. Scales were obtained by scraping thawed skins with a scalpel and then collected into a glass vial before drying overnight at 70 °C. Scales and otoliths were not processed further before submission for analysis of stable isotope contents. Tissue samples were sent to the University of Windsor GLIER Chemical Tracers Lab for isotopic analysis (Windsor, ON, Canada).

### Fin erosion

Assessment of fin erosion between rainbow trout and a wild salmonid of Parry Sound (lake trout *Salvelinus namaycush*) was also used as supporting evidence of the escape of rainbow trout from pens. Captive fish housed at high density show a high incidence of fin erosion (Person-Le Ruyet et al., 2007; Petersson et al., 2013). We compared damage to the caudal fins on photographs of rainbow trout and lake trout sampled in Parry Sound using available photographs of lake trout sampled for purposes other than the present study (e.g. Johnson et al., 2018). A scale of caudal fin damage adapted from Petersson et al. (2013) was established with the lower erosion level 1 (little to no erosion) what is typically seen in wild fish, intermediate erosion level 2 (clear erosion on less than 50% of the fin), and advanced erosion level 3 (fin more than 50% eroded) (Figure 1B).

### Statistics

Stable isotope data were submitted to a mixed-effect modeling analysis in Prism 8 (GraphPad Software, San Diego, CA), with tissue (liver, muscle, scales, otoliths) and source (captive, escaped) as fixed effects and individual fish as a random effect. Sidak’s multiple comparison test was used to assess the effect of source on each tissue. Analysis of covariance (ANCOVA) computed in SPSS Statistics 26 (IBM, Armonk, NY) was used to compare the relative size of brain and heart ventricle between captive and escaped trout. The same method was used for comparisons of relative brain size of trout captured by different methods or captured at different sites. Multivariate analysis of covariance (MANCOVA) in SPSS was used to evaluate the contribution of different regions to brain size differences between captive and escaped trout. Only trout in which all five brain regions could be measured accurately were included in this analysis. For both ANCOVA and MANCOVA, the body size variable was set as a covariate and all mass and length data were Log_10_ transformed to meet test assumptions.

## Results

### Evidence supporting escape from pens

Multiple lines of evidence support that the rainbow trout sampled in Parry Sound escaped from the pen culture operation shortly before harvest in fall 2018. First, all rainbow trout sampled outside growing pens were larger or close to market size (approx. 1 kg body mass), the size at which trout are reported to have escaped from growing pens (G. Cole, personal communication). Second, all rainbow trout caught were females, in line with the routine aquaculture practice of treating young fish to create monosexual growing stocks (Benfey, 1996). Third, our analysis of fin erosion showed that 75% of rainbow trout sampled in Parry Sound had intermediate or advanced fin erosion, a proportion similar to rainbow trout of similar size sampled directly from the pens (67%). Conversely, none of the 15 wild lake trout sampled in Parry Sound for which we have pictures available showed such fin damage. Fourth, no rainbow trout were captured during our fish sampling survey of Parry Sound in summer 2018 prior to the escape event, confirming the normal absence of this species from Parry Sound without input from the pen culture operation. Finally, we compared escaped and captive trout stable isotope signatures of carbon and nitrogen in tissues differing in molecular turnover rates in the fish sampled in 2019 (Figure 2). Results showed that liver δ^13^C signatures significantly differed between captive and escaped trout (Tissue*Source: F_(3, 88)_ = 3.5, p = 0.02; pen vs. escaped: *P* > 0.4 for otoliths, scales and muscle, *P* < 0.0001 for liver), with escaped trout showing more negative δ^13^C values suggesting an increased reliance on offshore food resources by the escaped trout (Vander Zanden and Rasmussen, 1999). Liver was the tissue with the fastest molecular turnover rate that we studied (Busst and Britton, 2018; Logan et al., 2006; MacNeil et al., 2006). The lack of difference in isotopic signatures in slower turnover tissues (muscle, scales and otoliths < liver) supports the common use of resources by all fish prior to escape (i.e. commercial fish feed). A lack of difference in liver δ^15^N signatures is likely due to comparable ^15^N content of fish feed and wild prey available to the escaped trout, which is supported by a comparison of δ^15^N in baseline organisms sampled from Parry Sound and commercial fish feed (Figure S1). The multiple lines of evidence presented above support our contention that rainbow trout sampled in the waters of Parry Sound had escaped from growing pens about 7 months prior to capture. The probability that some of the rainbow trout sampled in Parry Sound were strays from a nearby wild population is extremely low.

**Figure 2:**
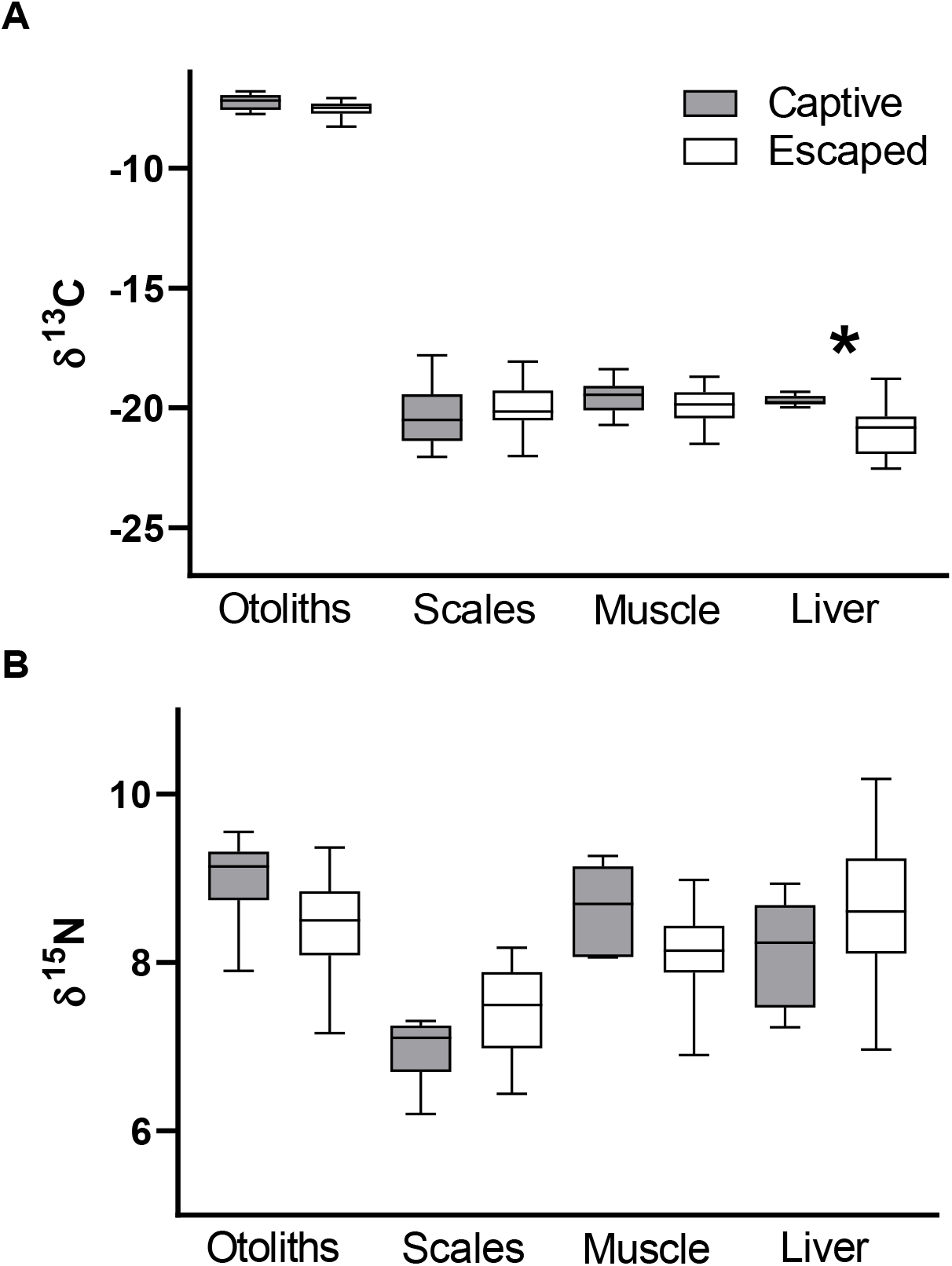
Comparison of stable isotope signatures in four tissues of captive and escaped rainbow trout. A) δ^13^Carbon, B) δ^15^Nitrogen. For each tissue and isotope, values for captive trout (gray bars) are compared to values for escaped trout (white bars). Boxes are medians and 25^th^ to 75^th^ percentiles, while whiskers are minimal and maximal values. The asterisk above liver values in panel A indicates a statistically significant Sidak’s multiple comparison test. Sample sizes for each tissue are 6 (captive) and 26 (escaped) trout sampled in 2019, except for scales of escaped trout (n=24).

### Brain size

We compared body size-brain size relationships of escaped trout and trout sampled directly from growing pens to test the prediction stating that increased complexity of life in a natural lake environment would increase brain size in escaped trout. Only trout above 330 mm fork length and 0.5 kg body mass were included in this analysis to ensure that the groups were within comparable size ranges. A preliminary analysis showed no difference in relative brain size of captive trout sampled in different years, so captive fish of different years were used as baseline for comparison with escaped trout. Figure 3A shows that brains of escaped trout are about 15% heavier on average than brains of fish captured directly from growing pens after accounting for body size. Importantly, relative brain size of most escaped trout fell above the range of variation seen within the captive trout sample, supporting a mechanism of brain size plasticity for the observed increase instead of selection against escaped trout with smaller brains. ANCOVA showed that the difference in brain size is statistically significant whether correction for body size is based on body mass (LogBodyMass: F_1_ = 166.5, *P* < 0.001, η_p_^2^ = 0.77; escaped vs. captive: F_1_ = 37.0, *P* < 0.001, η_p_^2^ = 0.43) or fork length (LogForkLength: F_1_ = 218.4, *P* < 0.001, η_p_^2^ = 0.82; escaped vs. captive: F_1_ = 7.8, *P* = 0.007, η_p_^2^ = 0.14). Inclusion of smaller captive trout collected in May 2019 in a supplementary analysis yielded similar results (LogBodyMass: F_1_ = 265.5, *P* < 0.001, η_p_^2^ = 0.81; escaped vs. captive: F_1_ = 31.4, *P* < 0.001, η_p_^2^ = 0.34), but these fish are excluded from Figure 3 for clarity.

**Figure 3:**
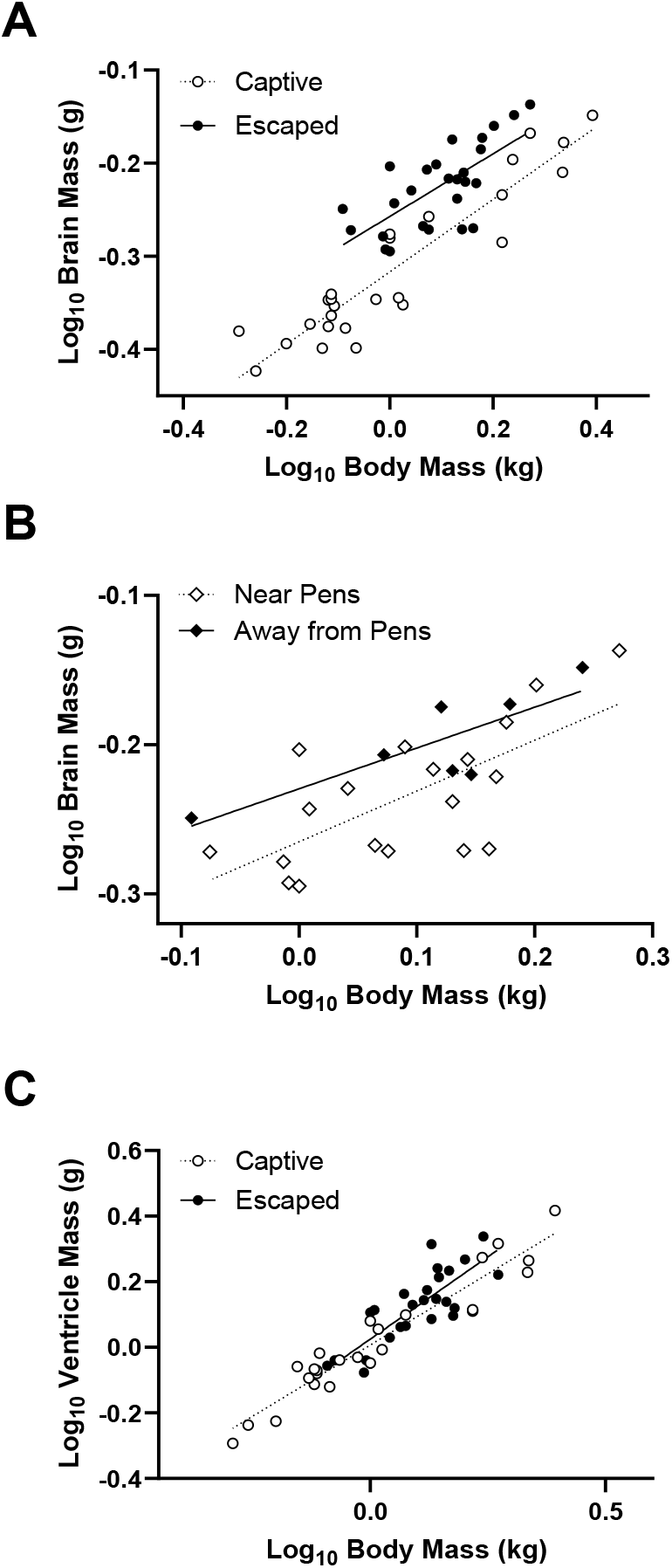
Effect of escape from a pen culture operation on brain size in rainbow trout. A) Body-brain mass relationships of captive trout sampled directly from growing pens (white symbols and dotted line) and escaped trout (black symbols and line). B) Body-brain mass relationships of escaped trout sampled in the vicinity of growing pens (white diamonds and dotted line) and escaped trout sampled far away from the pens (black diamonds and line). Relative brain size of escaped trout is larger than captive trout and larger in escaped trout captured farther away from the pen culture operation. C) Body-heart ventricle mass relationships of captive trout (white symbols and dotted line) and escaped trout (black symbols and line). There is no difference in relative heart ventricle size between captive and escaped trout. Linear regression was used to illustrate the relationships. Sample sizes are 26 captive (sampled in 2015: 9, 2016: 6, 2017: 5, 2019: 6) and 26 escaped trout (sampled in 2019) in panels A and C, and 19 near pens and 7 away from pens in panel B.

Since previous research established that larger brains relative to body size can facilitate the colonization of novel environments in birds and mammals (Fristoe et al., 2017; Sol et al., 2005; 2008), we were also interested in comparing brain size of escaped trout that moved away from the pen culture operation to those that remained in its vicinity. Even though we cannot ascertain the movements of trout during the 7 months following escape, local angling activity for escaped rainbow trout suggest that many fish remain in the vicinity of pens for an extended period. Therefore, capture at a great distance from the pens is at least an indicator that these fish dispersed away from the site of their escape and did not return near the pens daily. Figure 3B shows that trout captured in the northern part of Parry Sound in 2019, about 10 km due north from the pen culture operation, have larger brains than escaped trout captured near the pen culture operation. ANCOVA showed that this difference was on the statistical threshold (capture site: F_1_ = 4.3, *P* = 0.05, η_p_^2^ = 0.16) even though only 7 trout could be captured far away from the growing pens. This observation could support the notion that trout with the largest relative brain sizes were better suited to disperse in novel environments. This difference in brain size does not appear related to differences in foraging because liver stable isotope signatures do not differ between capture sites (Figure S2). It is also interesting to note that there is no relationship between relative brain size and liver stable isotope signatures among escaped trout (Figure S3A-B), suggesting no difference in diet based on brain size in escaped fish. Informal observation of stomach contents of the escaped trout captured in 2019 identified recently consumed prey as mostly littoral benthic macroinvertebrates (dragonfly and caddisfly larvae) and occasional forage fish.

Finally, we compared brain size of escaped trout captured by angling (n=10) and gill netting (n=16) to verify if angling pressure selectively removing smaller brained escaped trout could introduce a population bias contributing to the larger brains of escaped trout. ANCOVA showed no clear significant difference in brain size with capture method (F_1_ = 3.1, *P* = 0.09). The trend was for larger brains in trout captured by angling compared to trout captured by netting (ANCOVA EMM [95% CI]: angling, 0.62 [0.59-0.64]; netting, 0.58 [0.56-0.61]), which is opposite to how an angling bias could produce larger brains in the population of escaped trout.

### Brain region sizes

The size of five brain regions was measured to evaluate their contribution to the larger brain size observed in escaped trout. Figure 4 shows that the telencephalic brain regions located anteriorly (olfactory bulbs and rest of telencephalon) are generally larger in escaped trout. The relative sizes of the other brain regions overlap greatly between captive and escaped trout. MANOVA highlighted a statistically significant difference in region size between groups (LogBodyMass: F_5,34_ = 8.6, *P* < 0.001, η_p_^2^ = 0.56; escaped vs. captive: F_5,34_ = 9.5, *P* < 0.001, η_p_^2^ = 0.58). Follow-up univariate tests for each region showed that only the olfactory bulbs (F_1,38_ = 15.7, *P* < 0.001, η_p_^2^ = 0.29) and telencephalon (F_1,38_ = 36.3, *P* < 0.001, η_p_^2^ = 0.49) of escaped trout were larger compared to captive trout (about 36% and 40% larger, respectively). The other brain regions did not differ in size between groups (tectum: F_1,38_ = 2.2, *P* = 0.15, cerebellum: F_1,38_ = 0.04, *P* = 0.85, hypothalamus: F_1,38_ = 2.4, *P* = 0.13).

**Figure 4:**
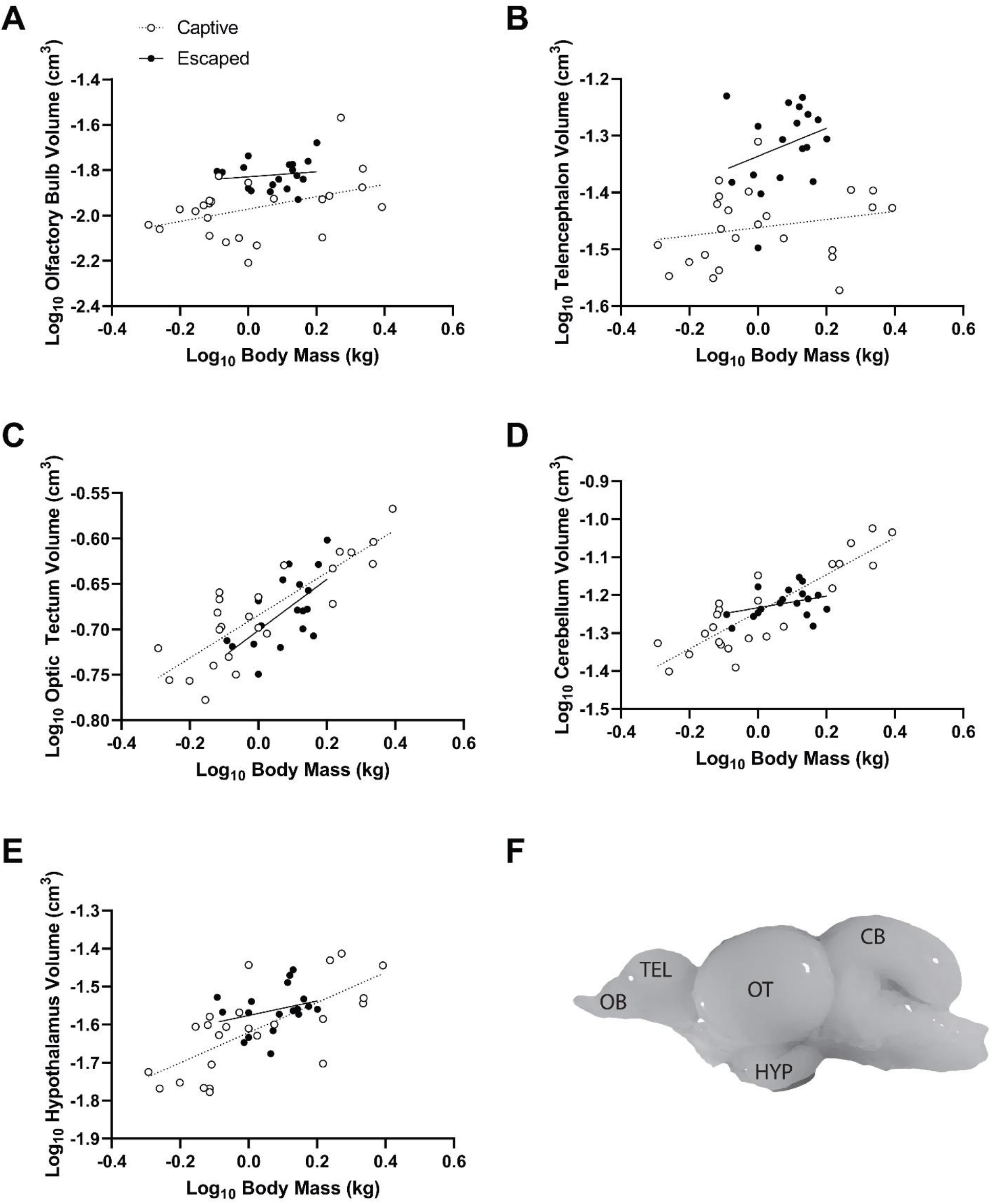
Effect of escape from a pen culture operation on brain region sizes in rainbow trout. Body-brain region volume relationships of captive trout sampled directly from growing pens (white symbols and dotted line) and escaped trout (black symbols and line). A) Olfactory bulb. B) Telencephalon. C) Optic tectum. D) Cerebellum. E) Hypothalamus. F) Lateral view of a rainbow trout brain illustrating the location of the five brain regions analyzed. Relative size of the olfactory bulbs and telencephalon is larger in escaped trout. Sample sizes are 24 captive and 18 escaped trout for all brain regions. Abbreviations: CB: cerebellum, HYP: hypothalamus, OB: olfactory bulb, OT: optic tectum, TEL: telencephalon.

### Heart ventricle size

To test our prediction that plastic changes for larger ventricles would be induced by the enhanced swimming requirements associated with life in a natural lake environment, we compared the heart ventricle size of escaped and captive trout. Preliminary analysis showed no difference in relative ventricle size of captive trout sampled in different years, but a more elongated ventricle shape of captive fish sampled in 2017 compared to other years. Therefore, we limited our analysis to relative heart size because year to year differences in early life conditions could have determined ventricle shape of the escaped trout (e.g. temperature differences: Dimitriadi et al., 2021). Figure 3C shows that the relationships between body mass and ventricle mass overlap greatly in escaped and captive trout. This observation is supported by a non-significant effect of escape on ventricle mass (LogBodyMass: F_1_ = 250.2, *P* < 0.001, η_p_^2^ = 0.84; escaped vs. captive: F_1_ = 2.7, *P* = 0.11). Thus, escape into the lake did not select for or induce the growth of larger heart ventricles.

## Discussion

While researchers have begun to recognize the plasticity of adult fish brain size from lab experiments, we used escaped aquaculture-raised rainbow trout to show the rapid change brain size can undergo when adult fish are newly exposed to a natural environment. Phenotypic flexibility of brain size is the best explanation for the observed difference between captive and escaped trout. As alternative explanations, the selective escape of large-brained trout can be ruled out because fall storms resulted in massive escape of tens of thousands of fish from broken pens without recovery. Secondly, selective removal of small-brained trout by angling can be rejected because angling capture showed no bias for small-brained trout. Finally, selective mortality of small-brained trout following escape is not supported by the data because relative brain size of most escaped trout was above the range of variation seen within the sample of captive trout. Escaped fish partitioned themselves into those that stayed near the aquaculture pens and those that moved away a long distance. Intriguingly, those that moved away had larger brains, possibly because they showed an even stronger increase in brain size in response to their novel wild environment, or because their larger brains promoted colonization of novel habitats (Fristoe et al., 2017; Sol et al., 2005; 2008). As rainbow trout went from a predictable schedule of pelleted feed in a simple, constrained floating pen environment to an expansive natural foraging arena where prey items were heterogeneous and evasive, we also expected rapid changes in the heart to aid with altered demands on locomotion. However, we found no difference in relative heart ventricle size between captive and escaped trout that would suggest differences in locomotion. Nevertheless, we found that stable isotope signatures in a fast turnover tissue of escaped trout showed a significant shift indicative of changing foraging conditions for increased open water feeding in escaped trout.

### Phenotypic flexibility of trout brain size

Our results contribute to mounting evidence showing that brain size in adult fish can be subject to phenotypic flexibility (see also Fong et al., 2019; Herczeg et al., 2015; Park et al., 2012; Turschwell and White, 2016). Flexibility of brain size would likely modulate cognitive capacity according to environmental complexity or foraging requirements, although the specific benefits of larger brains will require further investigation. Reducing brain size in a timely fashion is also likely advantageous in order to save resources for periods of high activity because nervous tissue is among the most energetically costly to maintain (Mink et al., 1981).

Flexibility of brain size associated with changing environmental conditions during lifetime is potentially widespread in organisms that maintain a high capacity for adult brain neurogenesis and lifelong brain growth, such as most anamniote and non-avian reptile vertebrates (Kaslin et al., 2008). Nonetheless, short-lived mammals living under constant high energy demands show seasonal cycles in skull and brain size that appear to match seasonal activity patterns (LaPoint et al., 2017; Lázaro et al., 2018; 2019). Further, seasonal and activity-dependent changes in regional size of the mammalian hippocampus and avian song control nuclei have been noted (Clayton and Krebs, 1994; Jacobs, 1996; Nottebohm, 1981; Tramontin and Brenowitz, 2000; Yaskin, 2011). These brain regions are characterized by abundant adult neurogenesis even though birds and mammals display overall determinate brain growth (Amrein et al., 2011; Goldman and Nottebohm, 1983). This suggests that the potential for phenotypic flexibility of brain size is not limited to basal vertebrates but is possibly limited to brain regions with high neurogenic potential. Despite the latter, differences in neurogenic potential across brain regions are unlikely to explain our finding that anterior telencephalic brain regions contributed most to the change in brain size observed in escaped trout because the brain region with the highest proliferative activity in teleosts appears to be the cerebellum (Zupanc and Horschke, 1995). Greater growth of telencephalic regions in escaped trout might be activity-dependent and reflect specific requirements of foraging involving olfactory and spatial processing, functions associated with the olfactory bulbs and dorsal telencephalon (Kotrschal et al., 1998; Rodríguez et al., 2002).

### Implications for studies of brain size evolution

The evolution of brain size has long attracted the interest of scientists (see Jerison, 1973). In studies of brain size evolution among taxa, researchers commonly use as little as one specimen to represent the ‘typical’ brain size of a given species (e.g. Clutton-Brock and Harvey, 1980; Garamszegi et al., 2002; Gonzalez-Voyer et al., 2009). Considering that brain size in many vertebrates may be subject to phenotypic flexibility, rapid plastic change within a lifetime could introduce important uncertainty in the ability to estimate brain size for a given species based on sampling conditions. We know little about the magnitude of plastic changes in brain size relative to differences that have evolved between species over evolutionary time, which could have an important impact on the evaluation of evolutionary patterns, especially at lower taxonomic levels. The average brain size difference between escaped and captive trout measured here provides an estimate of 15% in potential plastic change for this species. This means that using captive rainbow trout to establish the ‘typical’ brain size of this species would underestimate normal brain size by a substantial amount. Thus, establishing a species reaction norm of brain size should be considered, when possible, by estimating seasonal (e.g. McCallum et al., 2014), habitat (e.g. Axelrod et al., 2018) or other kinds of variation in relative brain size within a species and by factoring the captive or wild status of specimens. This variance around average brain size data could then be included in models of brain size evolution for more accurate evaluation of evolutionary patterns and their associated uncertainty.

### Relevance of organ phenotypic flexibility to fish-driven ecological dynamics

Evolutionary ecologists have long pushed the notion that rapid evolutionary responses have the potential to be major drivers of ecological dynamics (Hairston Jr et al., 2005; Thompson, 1998); a view that was later supported by experimental evidence (Yoshida et al., 2003). Despite this evidence, much of ecology research still ignores evolutionary dynamics as though they are too slow to significantly impact population dynamics (discussed in Endler, 1991; Thompson, 1998), perhaps because overall evidence from wild systems remains sparse (although see Turcotte et al., 2011). Here, we go beyond this growing literature by showing that a complex physiological structure (brain) can change on infra-evolutionary timescales in the wild. The role for brain size in fish cognitive capacity (Buechel et al., 2018) imply that change in this structure, or its trait distribution, can influence fish foraging capacity at the population level, which is a main determinant of fish effects on aquatic population dynamics. Plastic change in brain size has the potential to influence ecological dynamics directly or by interaction with heritable change (see Ellner et al., 2011). Therefore, top-down ecological dynamics in aquatic systems can be subject to drives at different time scales, from recurring periods in an individual lifetime to more or less rapid generational effects. The factors that determine which temporal drivers dominate under different conditions should prove fertile ground for future research.

### Can phenotypic flexibility contribute to ecosystem stability?

Ecologists have recently made arguments that higher order mobile predators can play major roles in mediating the stability of whole ecosystems if they can respond in a rapid and informed manner to spatial and temporal prey variation. Specifically, researchers have argued that if prey vary in multiple habitats non-synchronously then informed mobile predators can average across this variation like a stock market broker uses the “portfolio effect” across non-synchronous stocks to smooth variation over time and space providing stability in returns (McCann and Rooney, 2009; Schindler et al., 2015). Nonetheless, this mechanism requires that mobile organisms be capable of making rapid informed decisions, as delays in adaptive response to changing prey can drive significant instability (Abrams, 1992). Our results show that fish in the wild can indeed rapidly respond to novel environments by growing larger brains (15% growth) within a period of about 7 months. Therefore, it appears that fish have the physiological machinery to alter the ability to make informed decisions, as general theory for stability requires (e.g. McCann and Rooney, 2009), at a time scale faster than evolutionary mechanisms can provide. Thus, ecosystem stability mechanisms could also depend on cycles of energy budget management in long-lived predators (organ growth and shrinkage) that help smooth variance in cycles of population abundance over time. It remains to be seen if phenotypic flexibility of organs that contribute to foraging performance is a pronounced characteristic of mobile predators or a more widespread physiological phenomenon.

## Acknowledgements

We gratefully acknowledge Gord Cole and Kana Upton of Aqua Cage Fisheries and Cole-Munro (St-Thomas) who supplied us with farmed trout. James Simpson provided invaluable help in the field and Elizabeth Thurston helped in the laboratory. The Natural Sciences and Engineering Research Council of Canada Discovery Grant program (Laberge, McCann) and the Canada First Research Excellence Fund (McCann) provided financial support.

## Supporting Information Appendix

### Supplementary figures

**Figure S1:**
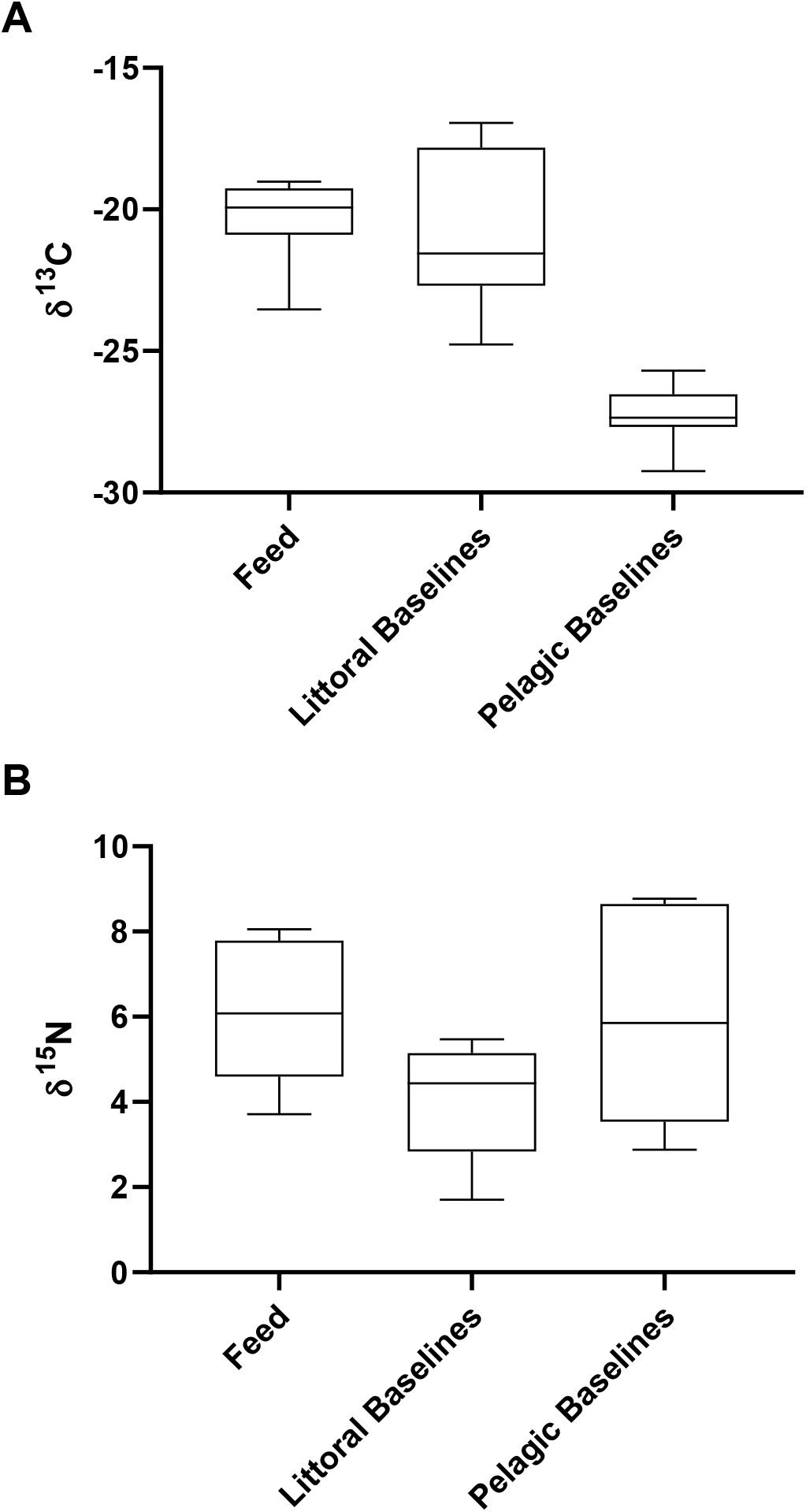
Comparison of stable isotope signatures of Parry Sound baseline organisms and commercial fish feed. A) δ^13^Carbon, B) δ^15^Nitrogen. Littoral baseline organisms were mayfly larvae (n=6) and snails (n=7), while pelagic baselines were zebra mussels (n=8). Feed was Skretting FW pellets 2.5-7.5 mm (n=5 samples) obtained in summer 2018. Boxes are medians and 25^th^ to 75^th^ percentiles, while whiskers are minimal and maximal values.

**Figure S2:**
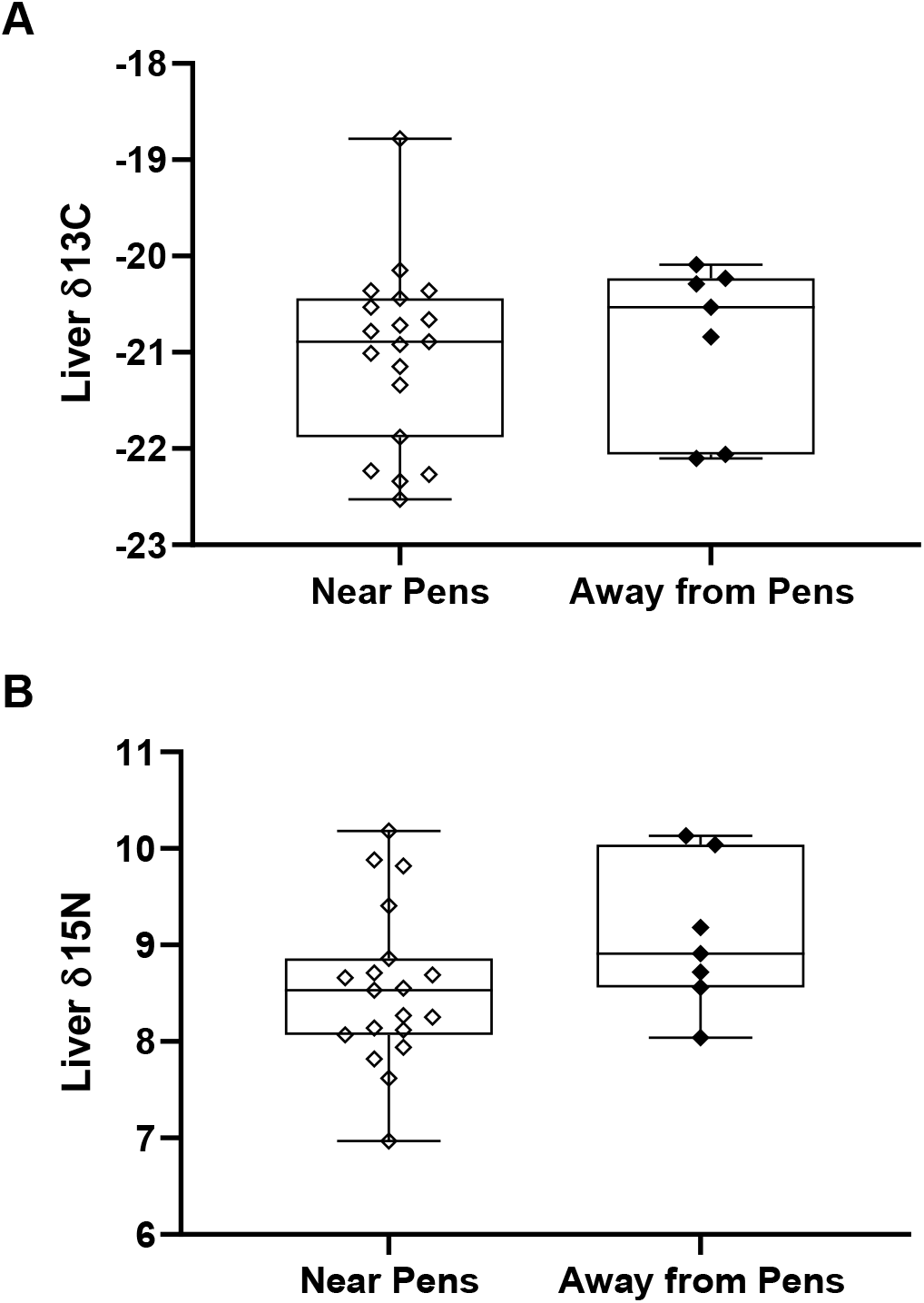
Liver stable isotope signatures and capture site of escaped trout. A) δ^13^Carbon, B) δ^15^Nitrogen. Boxes are medians and 25^th^ to 75^th^ percentiles, while whiskers are minimal and maximal values. Unpaired t-tests showed no difference in liver δ^13^C (t_24_ = 0.35, *P* = 0.73) or δ^15^N (t_24_ = 1.5, *P* = 0.15) by site of capture. Sample sizes are 19 (near pens) and 7 (away from pens) trout.

**Figure S3:**
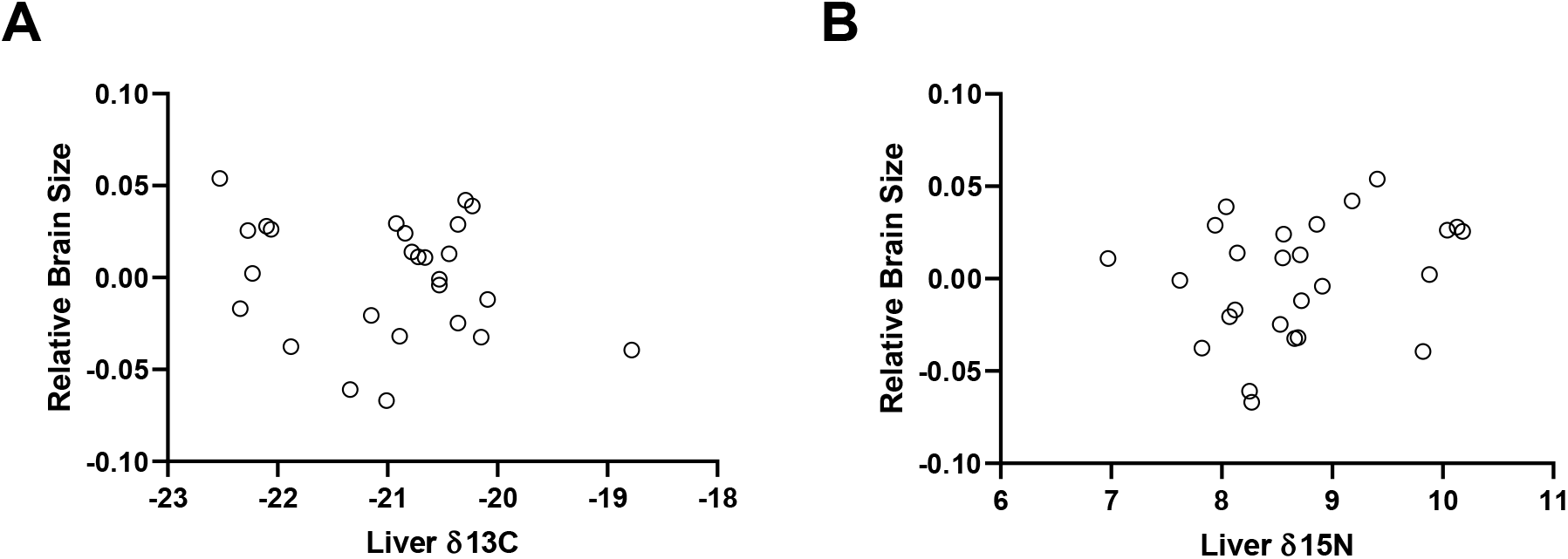
Absence of relationship between liver stable isotope signatures and brain size in escaped trout. Panels A (δ^13^Carbon) and B (δ^15^Nitrogen) show liver isotopic signatures in relation to relative brain size. Relative brain size is the residual values obtained from a linear regression of the logarithms of body mass and brain mass in escaped trout. Relationships between liver stable isotope signatures and brain size were assessed by linear regression: A: F_(1,24)_ = 0.98, *P* = 0.33; B: F_(1,24)_ = 1.78, *P* = 0.2. Sample size is 26 in both panels.

